# Modeling human population separation history using physically phased genomes

**DOI:** 10.1101/008367

**Authors:** Shiya Song, Elzbieta Sliwerska, Sarah Emery, Jeffrey M. Kidd

**Affiliations:** Department of Computational Medicine and Bioinformatics; Department of Human Genetics University of Michigan Medical School Ann Arbor, Michigan, USA

**Keywords:** Fosmid pool sequencing, population split-time, PSMC, MSMC, approximate Bayesian computation

## Abstract

Phased haplotype sequences are a key component in many population genetic analyses since variation in haplotypes reflects the action of recombination, selection, and changes in population size. In humans, haplotypes are typically estimated from unphased sequence or genotyping data using statistical models applied to large reference panels. To assess the importance of correct haplotype phase on population history inference, we performed fosmid pool sequencing and resolved phased haplotypes of five individuals from diverse African populations (including Yoruba, Esan, Gambia, Massai and Mende). We physically phased 98% of heterozygous SNPs into haplotype-resolved blocks, obtaining a block N50 of 1 Mbp. We combined these data with additional phased genomes from San, Mbuti, Gujarati and CEPH European populations and analyzed population size and separation history using the Pairwise Sequentially Markovian Coalescent (PSMC) and Multiple Sequentially Markovian Coalescent (MSMC) models. We find that statistically phased haplotypes yield an earlier split-time estimation compared with experimentally phased haplotypes. To better interpret patterns of cross-population coalescence, we implemented an approximate Bayesian computation (ABC) approach to estimate population split times and migration rates by fitting the distribution of coalescent times inferred between two haplotypes, one from each population, to a standard Isolation-with-Migration model. We inferred that the separation between hunter-gather populations and other populations happened around 120.0 to 140,000 years ago with gene flow continuing until 30,000 to 40,000 years ago; separation between west African and out of African populations happened around 70,000 to 80.0 years ago, while the separation between Massai and out of African populations happened around 50,000 years ago.

## Introduction

Haplotypes contain rich information about population history and are shaped by population size, natural selection, and recombination (VEERAMAH and HAMMER 2014; SCHRAIBER and AKEY 2015). Due to historic recombination events there are hundreds of thousands of pairs of loci along a chromosome that have distinct histories. Recent methodological advances permit the estimation of a detailed population demographic history from a single or several whole genome sequences based on the distribution of coalescent times across the genome. For example, Li and Durbin (LI and DURBIN 2011) developed the Pairwise Sequentially Markovian Coalescent model (PSMC) to reconstruct the distribution of the time since the most recent common ancestor (TMRCA) between the two alleles of an individual and infer population size changes over time. Typically, these TMRCA values are calculated using the two haploid genomes that compose the diploid genome of a single sample (LI and DURBIN 2011). When PSMC is applied to two haplotypes obtained from different populations, the inferred TMRCA distribution is informative about the timing of population splits since the time after which nearly no coalescence events occur is a good estimate for the population split time. One key question regarding human population history is the timing of population splits and the dynamics of separation between Africans and non-Africans, which has a great influence on modern genetic diversity. Li and Durbin (LI and DURBIN 2011) paired X chromosomes from African and non-African males and suggested that the two groups remained as one population until 60–80 kyrs ago with substantial genetic exchange up until 20–40 kyrs ago (assuming a mutation rate of 2.5 × 10^−8^ bp per generation and 25 years as generation time, estimates which approximately double when assuming a mutation rate 1.25× 10^−8^ bp per generation and 30 years as generation time (SCHIFFELS and DURBIN 2014)). Subsequently, PSMC applied to pseudo-diploid sequences was used to date the divergence time between non-human primate subspecies (PRADO-MARTINEZ *et al.* 2013). However, PSMC curves themselves provide only a qualitative measure of population separation and estimating split times is complicated by the presence of migration (PRITCHARD 2011).

MSMC (SCHIFFELS and DURBIN 2014) extends PSMC to multiple individuals, focusing on the first coalescence event for any pair of haplotypes. With multiple haplotypes from different populations, MSMC calculates the ratio between cross-population and within-population coalescence rates, termed the ‘relative cross coalescence rate’, a value reflecting population separation history. Schiffels et al. (SCHIFFELS and DURBIN 2014) applied MSMC on statistically phased genomes (two or four haplotypes per population) and suggested that African and non-African populations exhibited a slow, gradual separation beginning earlier than 200,000 years ago and lasting until about 40,000 years ago, while the median point of such divergence was around
60,0 – 80,000 years ago. The midpoint of the relative cross-coalescence decay curve has been used as an estimate of population separation time (TEWHEY *et al.* 2011; SCHIFFELS and DURBIN 2014). Although useful, this approach does not generate parametric estimates for population history under standard models. As none of these methods to infer population separation history were applied on physically phased genomes, it is unclear how phasing errors and missing data affect this type of analysis.

In this manuscript, we construct physically phased genomes of five individuals from diverse African populations (including Yoruba, Esan, Gambia, Massai and Mende). We reanalyzed fosmid sequencing data for individuals from the Gujarati, San and Mbuti populations and assess the ability to correctly assemble SNP haplotypes using fosmid pool sequencing and compare the resulting data with statistically phased haplotypes. We have previously compared several reconstructed haplotypes from a subset of these samples with those released by 1000 Genomes Phase III Project (CONSORTIUM 2015). In this paper, we focus on how well the existing statistical phasing software SHAPEIT (DELANEAU *et al.* 2012) performs given the available 1000 Genomes reference panel and how different reference panels perform, especially for samples from populations not represented in the panel. We further assess the impact of phasing error on MSMC’s estimates of population split times using physically phased genomes vs. statistically phased genomes. Finally, we extend the current PSMC method to model population splits. We apply an approximate Bayesian computation (ABC) method to obtain posterior estimates of split time and migration rate by fitting the inferred TMRCA distribution obtained from PSMC on pseudo-diploid genomes to a standard Isolation-with-Migration model. Additionally, we assess the sensitivity of existing methods to missing data and phasing errors from statistically phased haplotypes.

## Materials and Methods

### Reconstructing Haplotypes Using Fosmid Pool Sequencing

We performed fosmid pool sequencing and standard Illumina sequencing on individuals NA19240, HG03428, HG02799, HG03108, and NA21302, the detailed methods of which is elaborated in the 1000 Genomes Phase3 paper (CONSORTIUM 2015) (Supplemental Table 1–(3). Paired end reads were aligned to the reference genome assembly (GRCh37, with the pseudoautosomal regions of the Y chromosome masked to ‘N’) using BWA v0.5.9−r16 (LI and DURBIN 2009). PCR duplicates were removed by Picard v1.62. Reads in regions with known indels were locally realigned and base quality scores were recalibrated using GATK (MCKENNA *et al.* 2010). We generated GVCF files (Genomic VCF) with a record for every position in the genome using GATK HaplotypeCaller v3.2–2. Variants were called using GenotypeGVCFs and filtered by applying Variant Quality Score Recalibration(VQSR) implemented in GATK to select a SNP set that included 99% of sites that intersect with the HapMap and 1000 Genomes training set. We define callable regions as sites that are within half and 2 times the average coverage and with genotype and mapping quality scores greater than 20. We kept variants that either passed VQSR filtering or were present in the 1000 Genomes Phase I reference panel, which served as the starting point for subsequent haplotype phasing. We followed the procedure described in 1000 Genomes Phase3 paper (CONSORTIUM 2015) to process fosmid sequencing data. We generated phased haplotypes from five individuals plus samples NA20847 (KITZMAN *et al.* 2011), HGDP01029 and HGDP00456 (MEYER *et al.* 2012) that were published previously and obtained haplotype phased blocks using ReFHap (DUITAMA *et al.* 2012).

For NA19240, HG02799, HG03108 and NA21302, we used phase-determined SNPs from trio genotyping available from HapMap and AffyMetrix to guide paternal and maternal allele assignment within blocks. We determined paternal and maternal allele identity based on the majority of phased SNP assignments, then identified and corrected corrected switch errors only if the increase in MEC value (minimum error correction) was less than 50 after correction. For NA20847, HG03428, HGDP01029, and HGDP00456, phase-determined SNPs from trio data are unavailable. For these samples, we applied Prism (KULESHOV *et al.* 2014), a statistical phasing algorithm designed to assemble local blocks into long global haplotype contigs. This method is an extension to the Li and Stephens‘s model (LI and STEPHENS 2003) that utilizes a reference panel of phased haplotypes and a genetic map of the genome with an additional parameter representing the phase of each block in the hidden Markov model to enforce the locally phased structure at the global phasing level. We grouped local blocks into windows with size smaller than 1Mbp and with at least 2 local blocks. Each window overlapped by 1 local block, which was used to link adjacent windows together. For sample NA12878, we directly used the phased SNP haplotypes constructed by fosmid pool sequencing from a previous study (DUITAMA *et al.* 2012), downloaded from http://www.molgen.mpg.de/~genetic-variation/SIH/data, and we obtained callable regions and high-confidence SNP call sets from the sequencing results of 1000 Genomes Pilot Project (CONSORTIUM 2010) to construct full haplotypes.

### MSMC analysis

We applied the Multiple sequentially Markovian coalescent (MSMC) (SCHIFFELS and DURBIN 2014) model on four haplotypes, two haplotypes per individual each population. We used ‘fixedRecombination’ and ‘skipAmbigous’ for inference of population separation. MSMC analysis yields inferred cross-population and within-population coalescence rates. We calculated the relative cross coalescence rate (RCCR) by dividing the cross-population coalescence rate by the average of within-population coalescence rate and plotted it as a function of time. We also applied MSMC on individual diploid genomes, which is very similar to PSMC, with subtle differences due to the underlying model SMC’ (MARJORAM and WALL 2006) versus SMC (MCVEAN and CARDIN 2005). In order to differentiate it from PSMC, we refer to such analysis as PSMC’.

### PSMC on pseudo-diploid genome

Pairwise sequentially Markovian coalescent (PSMC) (LI and DURBIN 2011) inference was performed as previously described (LI and DURBIN 2011). PSMC builds a HMM to infer the local TMRCA based on the local density of heterozygotes. In the model, hidden states are discretized TMRCA values, and transitions represents ancestral recombination events. On autosomal data, we use the default setting with T_max_=15, n=64, and pattern ‘1*4+25*2+1*4+1*6’. When applying PSMC on a pseudo-diploid genome, there are four possible configurations of the two haplotypes, namely hap1-hap1, hap1-hap2, hap2-hap1, hap2-hap2. We applied PSMC to each possible configuration and took the average of the estimates. We obtained the inferred TMRCA distribution directly from PSMC output, the fifth column representing the fraction of the genome that coalesced in an indicated TMRCA bin.

### ABC analysis

We implemented an ABC framework to estimate split time and migration rate given the inferred TMRCA distribution from PSMC output. We computed the coalescence time density of two chromosomes based on the Isolation-With-Migration model (WANG and HEY 2010; HOBOLTH *et al.* 2011) and integrated coalescence time density on the 64 time intervals in which PSMC is parameterized. We use chi-square statistics calculated between the observed TMRCA distribution obtained from PSMC output and the computed one as the distance between estimates in the ABC framework.

We formulate the IM model as continuous time Markov chain (WANG and HEY 2010; HOBOLTH *et al.* 2011). The rate matrix Q is given by: 
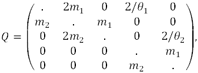
where the states are S_11_(both gene are in population 1), S_12_ (one gene is in population 1 and the other is in population 2), S_22_ (both gene are in population 2), S_1_ (the genes have coalesced and the single gene is in population 1), S_2_ (the genes have coalesced and the single gene is in population 1), and *θ_1_* and *θ_2_* is the scaled population sizes, and m_1_ and m_2_ are the migration rates. The density of coalescence time can be calculated as follows: 
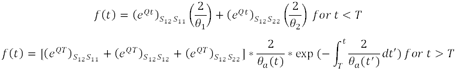
where T is the split time and θ_*a*_(t) the ancestral population size. We use the ancestral population size inferred from PSMC of the pseudo-diploid genome as the ancestral population size, and use the inferred population size of each diploid genome (from PSMC) as the population size for each population after the split. For African populations, we assume constant population size after the split. For non-African populations, we assume that the population experienced a bottleneck event after the split and experienced population growth beginning 40 kyrs ago. For our ABC framework, the parameters of interests are T (split time) and m (migration rate after the split). We assumed a uniform prior for the split time and time when migration ends, and a uniform prior on migration rate in log10 scale, and applied an ABC method based on sequential Monte Carlo (TONI *et al.* 2009) (SMC) to the parameter estimation, since it can be easily run in parallel and is more efficient than an ABC rejection sampler. We drew a pool of 5000 candidate parameter values (called particles) from the prior distribution. Instead of setting the final stringent cut-off ε (if the distance between summary statistics are lower than ε, we accept it), we gradually lowered the tolerance ε_1_ > ε_2_ >ε_3_ >> 0, thus the distributions gradually evolve toward the target posterior. The first pool generated by sampling from the prior distribution. The particles that were accepted using the first threshold >_1_ were sampled by their weights and perturbed to get new particles. As the tolerance threshold lowered to the final cut-off, we obtained the target posterior distribution. In each iteration, we choose the threshold ε such that 20% of particles are accepted, achieving N=1000 accepted particles. The perturbation kernels for all parameters are uniform, K=σU(−1,1), with σ equal to 20% of the difference between maximum and minimum values. We perform three iterations and summarized the mean, median and 95% HPD confidence interval for each parameter. For simulations, we generated 100 30Mb sequences of two individuals representing African and European samples and having split times ranging from 60 kyrs to 150 kyrs ago, with subsequent migration until 30 kyrs ago using MaCS (CHEN *et al.* 2009).

## Results

### 1. Haplotype reconstruction

We utilized RefHap to reconstruct haplotypes using fosmid pool sequencing of 8 individuals from diverse populations (NA19240 (Yoruba), HG02799 (Gambia), HG03108 (Esan), HG03428 (Mende), NA21302 (Maasai), HGDP01029 (San), HGDP00456 (Mbuti) and NA20847 (Gujarati) and obtained phased haplotypes for NA12878 (CEU) from a previous study (DUITAMA *et al.* 2012; PRÜFER *et al.* 2014). In total across all pools, each genome was covered by an average of 6–17 clones with a median sequence coverage ranging 16.9–24.8x (Supplemental Table 2). The effect of increased clone counts on phased block size is dramatic: when doubling the number of fosmid clones, the N_50_ of phased blocks tripled, achieving over 1Mbp for four of the African samples (Figure 1, Table 1).

**Figure 1.**
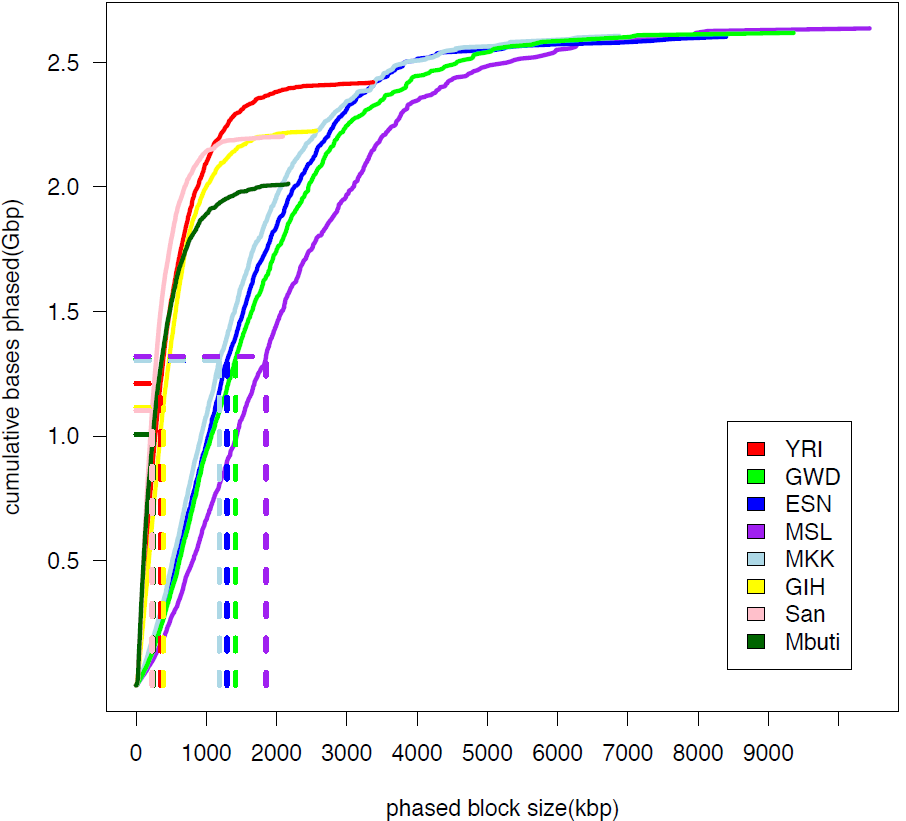
**Haplotype assembly results**. The relationship of block size and the cumulative length of constructed haplotypes are plotted. Dashed lines indicate the N50 of phased blocks, the block length such that 50% of the total is represented by blocks of that size or greater.

Although SNPs within each block are phased, the relationships between blocks cannot be directly established due to the absence of linking fosmid clones. We utilized two approaches to overcome this limitation. For samples that are members of genotyped trios, we utilized SNP transmission patterns to link adjacent blocks together producing near-to-complete haplotypes, encompassing over 97% of total heterozygous SNPs for HG02799, HG03108, NA21302, and 92.7% for NA19240. Comparison with deterministically phased SNPs identified potential switch errors due to insufficient clone support within our inferred haplotypes, which we corrected prior to subsequent analysis (Table 1, Supplemental Figure 1). We find 99.66% concordance between the fosmid-phased SNPs for NA19240 and heterozygous SNPs phased based on transmission from this sequenced trio (CONSORTIUM 2010). We further compared our phased haplotypes for NA19240 to the sequence of 33 fosmid clones from the same individual (KIDD *et al.* 2008), observing differences at 5 of the 1,013 heterozygous sites (0.5%) encompassed by the 33 clones (Supplemental Table 4). In total, the aligned clones encompass 1,102,213 bp excluding alignment gaps, and have 51 single nucleotide differences in comparison with our data. If we assume that all of these differences are errors in our inferred sequences, this suggests that our haplotypes have an overall sequence error rate of less than 0.005% or a Phred (EWING and GREEN 1998) quality score greater than Q40.

**Table 1.** **Phasing statistics from fosmid pool sequencing**. We resolved haplotypes using fosmid pool sequencing. MEC is the number of entries to correct when resolving haplotypes. Switch errors are counted as the number of switches required to obtain the same haplotype phase when comparing inferred haplotype phase with true haplotype phase. Switch error rate is switch error normalized by number of variants for comparison. For samples labeled with *, we applied Prism to link adjacent block together

| Populat<br>ion | Sample | Number of<br>Clones<br>After Filter | Number<br>of<br>Blocks | N50<br>(kbp) | MEC<br>value | Number of<br>SNPs to be<br>phased | % Phased<br>SNPs<br>within<br>blocks | **Number**<br>**of** blocks<br>assigned<br>parental<br>allele | Percent<br>SNPs<br>assigned<br>parental<br>allele | Switch<br>Error<br>Count | Switch<br>Error<br>Corrected | Switch<br>Error<br>Rate |
| --- | --- | --- | --- | --- | --- | --- | --- | --- | --- | --- | --- | --- |
| YRI | NA19240 | 521,783 | 16,334 | 347 | 37,143 | 2,588,454 | 92.94% | 15,171 | 92.74% | 586 | 421 | 0.091% |
| GWD | HG02799 | 1,141,020 | 5,236 | 1416 | 82,387 | 2,780,269 | 99.00% | 3,041 | 98.38% | 327 | 146 | 0.163% |
| ESN | HG03108 | 1,058,027 | 5,416 | 1294 | 77,499 | 2,756,725 | 99.07% | 3,323 | 98.45% | 258 | 115 | 0.127% |
| MKK | NA21302 | 892,863 | 6,097 | 1416 | 175,935 | 2,736,727 | 98.60% | 3,751 | 97.96% | 336 | 265 | 0.098% |
| MSL | HG03428* | 1,424,234 | 4,390 | 1849 | 167,294 | 2,775,099 | 99.30% | 3,549 | 97.90% | - | - | - |
| GIH | NA20847* | 571,419 | 16,838 | 385 | 44,870 | 1,680,704 | 93.37% | 13,319 | 90.98% | - | - | - |
| San | HGDP01029* | 358,759 | 17,695 | 228 | 27,712 | 2,623,001 | 87.10% | 16,516 | 89.15% | - | - | - |
| Mbuti | HGDP00456* | 381,075 | 18,465 | 242 | 28,629 | 2,517,569 | 78.70% | 17,385 | 81.72% | - | - | - |

**Table 2.** **Comparison between statistical phasing**. We calculated haplotype concordance, switch error rate, flip error rate, mean inter-switch distance, and mean length of incorrectly phased haplotype between haplotypes resolved by fosmid pool sequencing and haplotypes statistically phased using either the 1000 Genomes Phase1 or Phase3 reference panels. * indicates that trio data was unavailable to link blocks together and phasing comparison analysis was limited to comparisons within RefHap blocks.

| Individual | Haplotype Concordance | Switch Error Rate | Flip Error Rate | Mean inter-switch distance(kbp) | Mean length of incorrectly phased haplotype(kbp) |
| --- | --- | --- | --- | --- | --- |
| fosmid phased haplotypes vs SHAPEIT phased haplotypes using 1000 Genomes Phase1 reference Panel |  |  |  |  |  |
| NA19240 | 54.60% | 1.33% | 0.60% | 84.6 | 69.6 |
| HG02799 | 52.46% | 1.84% | 0.79% | 52.2 | 43.3 |
| HG03108 | 53.62% | 1.05% | 0.47% | 94.1 | 78.8 |
| NA12878 | 53.18% | 0.87% | 0.32% | 170.0 | 144 |
| NA21302 | 52.00% | 2.32% | 1.02% | 43.6 | 37.6 |
| HG03428* | 70.01% | 1.88% | 0.95% | 42.6 | 28.5 |
| NA20847* | 79.30% | 1.83% | 0.97% | 46.5 | 29.5 |
| HGDP01029* | 69.83% | 6.87% | 3.50% | 12.5 | 7.3 |
| HGDP00456* | 78.09% | 4.68% | 2.70% | 16.1 | 8.8 |
| **Average** | **62.57%** | **2.52%** | **1.26%** | **62.5** | **49.7** |
| fosmid phased haplotypes vs SHAPEIT phased haplotypes using 1000 Genomes Phase3 reference Panel |  |  |  |  |  |
| NA19240 | 68.00% | 0.33% | 0.21% | 480.5 | 293.6 |
| HG02799 | 77.10% | 0.63% | 0.27% | 296.5 | 124.4 |
| HG03108 | 69.40% | 0.42% | 0.27% | 346.5 | 208.5 |
| NA12878 | 58.90% | 0.67% | 0.32% | 264.4 | 204.4 |
| NA21302 | 53.10% | 2.44% | 1.08% | 41.2 | 32.9 |
| HG03428* | 89.70% | 0.66% | 0.50% | 132.2 | 56.1 |
| NA20847* | 91.50% | 1.00% | 0.73% | 70 | 36.9 |
| HGDP01029* | 69.97% | 7.17% | 3.77% | 12 | 6.9 |
| HGDP00456* | 77.47% | 5.08% | 2.97% | 14.9 | 8.1 |
| **Average** | **72.79%** | **2.04%** | **1.12%** | **184.2** | **108.0** |
| Fosmid phased haplotypes: assign parental alleles using trio data vs using Prism |  |  |  |  |  |
| NA19240 | 54.12% | 0.05% | 0.00% | 1242.6 | 1115.0 |
| HG02799 | 58.18% | 0.02% | 0.00% | 3427.3 | 2821.9 |
| **Average** | **56.15%** | **0.03%** | **0.00%** | **2335.0** | **1968.5** |

For individuals HG03428 (MSL), NA20847 (GIH), HGDP01029 (San), and HGDP00456 (Mbuti), trio data is unavailable. For these samples, we assigned 80%–98% of SNPs to a parental allele using Prism (KULESHOV *et al.* 2014), a statistical phasing algorithm designed to assemble short local blocks into longer global haplotype contigs. To evaluate how well Prism performs in this context of large haplotype-block assignment, we applied Prism to NA19240 and HG02799 and compared the assignment of local blocks with our assignment based on trio phase-determined SNPs. For NA19240, 6575 out of 13591 blocks (47.6%) were assigned differently, affecting 45.88% of total heterozygous SNPs. For HG02799, 1214 out of 2810 blocks (43.2%) were assigned differently, affecting 41.82% of total heterozygous SNPs. This results in mean interswitch distance of 2335 kbp and mean incorrectly phased haplotype length of 1967 kbp, with a 0.03% switch error rate.

### Comparison with statistical phasing

We applied SHAPEIT (DELANEAU *et al.* 2012) using either the 1000 Genomes Phase 1 reference panel(ABECASIS *et al.* 2012) (1092 individuals, 14 populations) or Phase3 reference panel(CONSORTIUM 2015) (2504 individuals, 27 populations) separately to statistically phase each individual (Supplemental Figure 2, Table 2). For haplotypes phased using the 1000 Genomes Phase 1 reference panel, the average switch error rate is 2.52%, half of which are flip errors, namely single alleles appearing on the opposite haplotype. Haplotypes phased using the Phase3 reference panel have a higher concordance rate (72.79%), longer mean length of incorrectly phased haplotype (108.0 kbp) and mean inter-switch distance (184.2 kbp), but similar levels of switch error rate (2.04%) and flip error rate (1.12%). This reflects the high accuracy of the 1000 Genomes Phase 3 release haplotypes, a result of a multi-stage phasing process that utilized a haplotype scaffold of trio-genotyped SNPs. For NA21302, HGDP01029, HGDP00456, whose populations are not included in 1000 Genomes reference panel, the level of switch errors and incorrectly phased haplotype were similar using either the Phase1 or Phase3 reference panel. For HG03428, NA20847, HGDP01029 and HDP00456, the comparisons of haplotypes are restrained to within blocks since blocks were statistically linked into long global haplotypes.

### The impact of phasing error on inference using MSMC

We applied PSMC’, similar to PSMC but using the SMC’ framework to perform demographic inference on nine individuals from nine populations. We assumed a human mutation rate of 1.25×10“^8^ bp per generation and 30 years as generation time, although results can be easily rescaled for comparison with other estimates (KONG *et al.* 2012; SCHIFFELS and DURBIN 2014). Consistent with previous findings, the PSMC’ curves of the nine individuals revealed that all populations shared the same two-fold increase of ancestral population size prior to 300 kyrs ago, after which the inferred population size of the African populations began to differentiate from non-African populations with all gradually experiencing an effective population size reduction (Figure 2), although we note that the simulations indicate that such shifts in PSMC curves may overestimate the timing of population size changes (LI and DURBIN 2011; PRUFER *et al.* 2014; HENN *et al.* 2015). Non-African populations experienced a ten-fold reduction of effective population size, but experienced a rapid population growth after 30 kyrs ago. Such observations were equivalent to previous PSMC analysis on diploid genomes after adjusting for differences in assumed mutation rate (LI and DURBIN 2011; MEYER *et al.* 2012; SCHIFFELS and DURBIN 2014).

**Figure 2.**
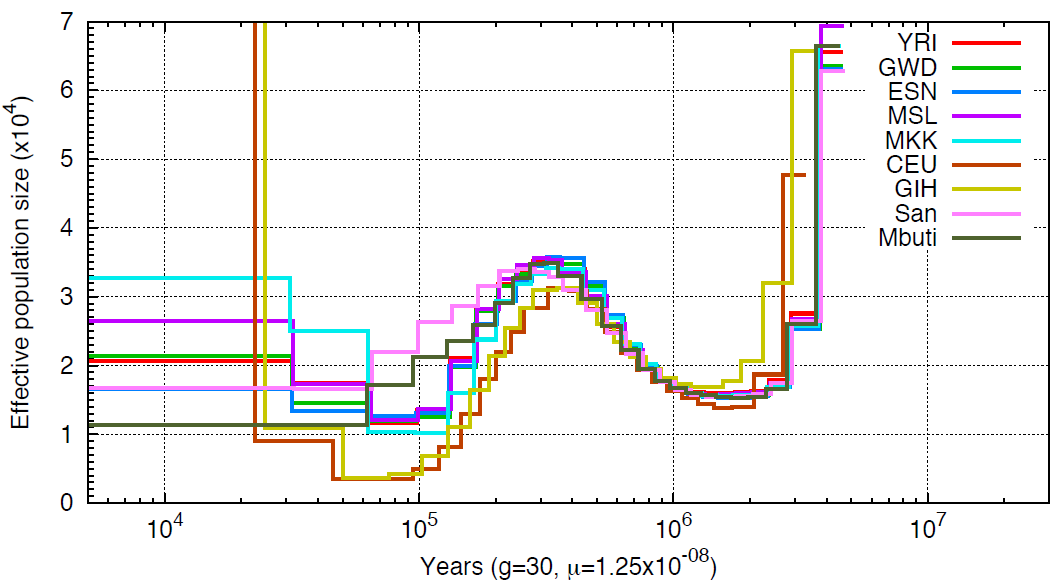
**PSMC’ inferred population history**. Population sizes inferred from the autosomes of nine individuals from nine populations are shown.

The multiple sequentially Markovian coalescent (MSMC) (SCHIFFELS and DURBIN 2014) model extends PSMC to multiple individuals. MSMC estimates the relative cross population coalescence rate, which drops from one to zero as populations separate. We applied the Multiple sequentially Markovian coalescent (MSMC) model on four physically phased haplotypes, two haplotypes per individual from each population and plotted the relative cross coalescence rate as a function of time (Supplemental Figure 3), using the time when RCCR drops to 50% as an estimate of the split time (Figure 3). We noticed that the more ancient the split event, the wider the inferred time interval. A similar pattern was also observed using simulated data (Supplemental Figure 4). We also performed MSMC analysis on haplotypes inferred using SHAPEIT (triangle, Figure 3). Statistically phased haplotypes show a more recent separation time and a narrower time span, particularly for comparisons involving San or Mbuti samples.

**Figure 3.**
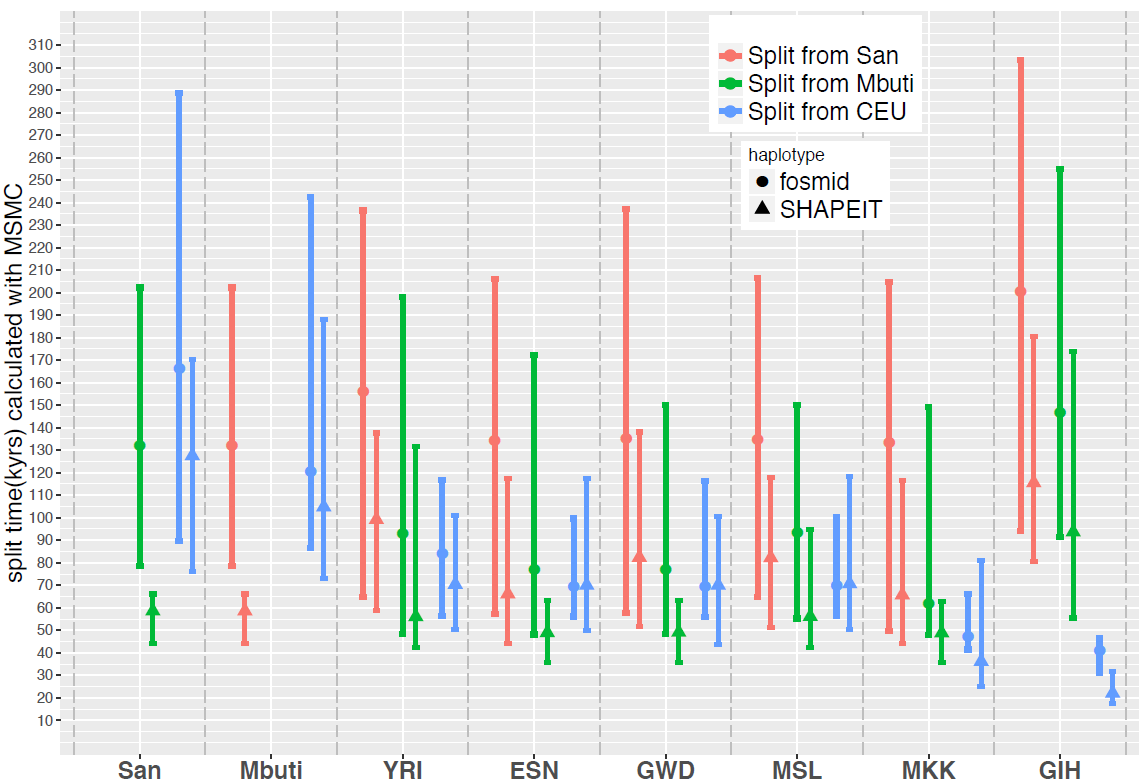
**MSMC inferred split times**. Circles or triangles represent the time when the cross coalescence rate dropped to 50%, with lines representing the time when cross-coalescence rate reached 25% and 75%. Inferred split times were inferred using haplotypes phased by the fosmid pools approach (circle) or SHAPEIT (triangle).

### An ABC method to infer population split time using PSMC on pseudo-diploid genomes

PSMC applied to pseudo-diploid samples also provides information on population separation history. If population splits are total and sudden, no coalescent events between populations will occur after their separation. Thus, when applying PSMC on a pseudo-diploid individual where one chromosome comes from one population and the second chromosome comes from another population, the time when the PSMC estimate of N_e_ goes to infinity provides an estimate for the population split time (LI and DURBIN 2011). However, the inferred PSMC curve usually increases in a step-wise manner, making it difficult to determine the exact time of split event. Subsequent migration after the split is a further confounding factor (PRITCHARD 2011).

To better interpret pseudo-diploid PSMC curves (Figure 4 and Supplemental Figure 5), we implemented an ABC framework to estimate the population split time and migration rate given the TMRCA distribution inferred from the PSMC output. We compared the observed TMRCA distribution with the analytical distribution determined by an Isolation-With-Migration model(WANG and HEY 2010; HOBOLTH *et al.* 2011) with the indicated values for split time and post-separation migration and applied an ABC method based on sequential Monte Carlo(TONI *et al.* 2009) (also abbreviated as ABC-SMC) to estimate the target posterior distribution of each parameter. We tested this approach using simulated data with a split time ranging from 60 kyrs to 150 kyrs ago, with subsequent migration continuing until 30 kyrs ago (Supplemental Figure 6). >For each split-time, we considered three levels of symmetrical migration: 2×10^−5^, 10×10^−5^, 40×10^−5^. For small levels of migration, the inferred split is quite accurate, with the mean value of the posterior distribution centered on the true value. However, for larger migration rates the inferred split-time tends to be smaller than the true value. This bias is exacerbated with subsequent iterations of ABC sampling. The magnitude of the inferred migration rate is reasonably accurate, as observed in the log10 scale.

**Figure 4.**
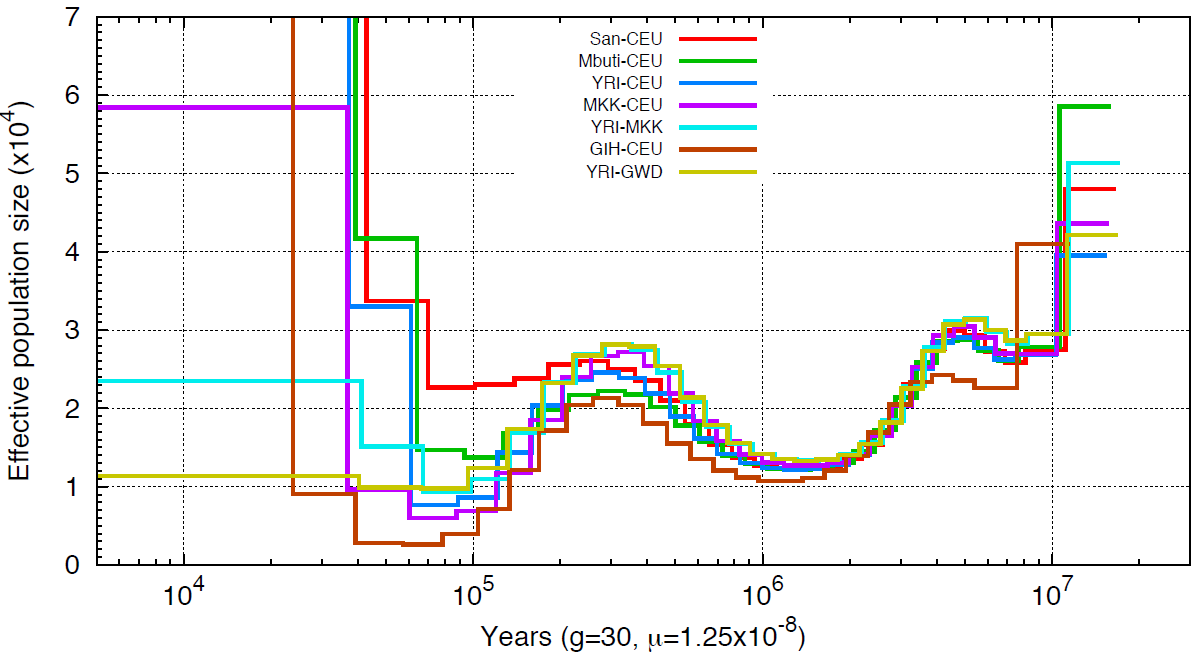
**PSMC on pseudo-diploid genomes**. Population sizes inferred from combined autosomes, with one haplotype chosen from each population are shown. Plotted curves are the average results obtained from four possible global haplotype configuration, namely hap1- hap1,hap1-hap2,hap2-hap1,hap2-hap2. Haplotypes were constructed using the fosmid pool approach.

An additional complication in the application of this method to real data is the treatment of unphased sites, which generally impact less than 10% of SNPs in each comparison (Supplemental Table 5). Using our simulations, we evaluated three methods for processing unphased SNPs: 1) randomly assigning the phase, 2) marking unphased sites along with all homozygous segments ending in an unphased heterozygous site as missing data (as recommended for MSMC) (SCHIFFELS and DURBIN 2014) and 3) marking only unphased SNPs as missing data. Even with 10% of unphased sites, the third method results in a PSMC curve similar to the original, while the first two methods give PSMC curves shifted to an earlier increased effective population size, which may result in an earlier inferred split time (Supplemental Figure 7). We therefore applied the third method to unphased SNPs in our analysis.

### Inferred split times using physically phased genomes

We applied our ABC method to date the split-times among African and European populations (Figure 5, Supplemental Figure 8). We find that the San population separated from the other samples the earliest, around 120 kyrs to 140 kyrs ago, with subsequent migration rate around 10~15×10^−5^ until 30–40 kyrs ago, an estimate that is more recent than that obtained from MSMC analysis (the median point of divergence using MSMC of San from other African populations was around 130 kyrs ago, and 160 kyrs ago with CEU population). The separation between west African and CEU populations occurred 70–80 kyrs ago with migration at a rate of 8~40×10^−5^ until 30 kyrs ago, while Maasai separated from the CEU population around 50 kyrs ago with a greater amount of gene flow until present, with migration rate on the magnitude of 10^−3^.

**Figure 5.**
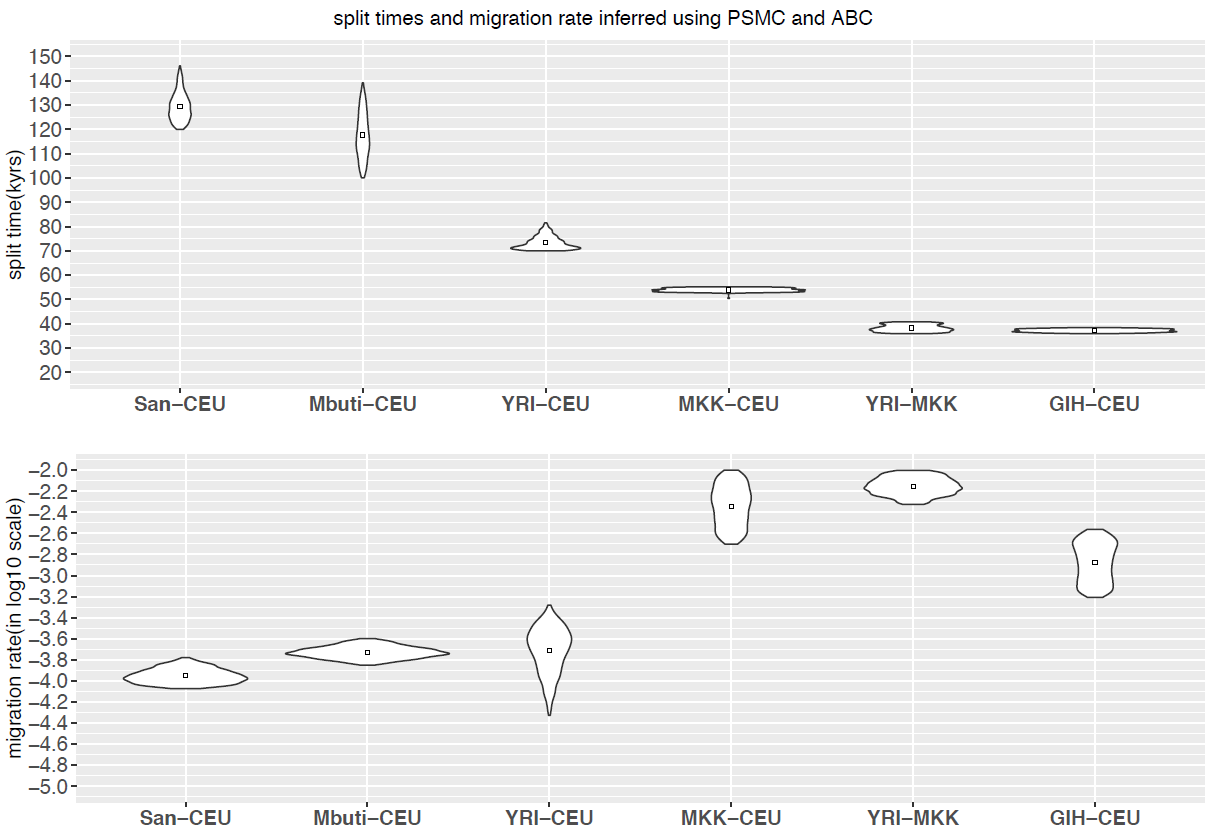
**Split times and migration rate inferred using PSMC and ABC**. We implemented ABC-SMC framework to estimate split time (A) and migration rate (B) given the inferred TMRCA distribution obtained from PSMC output. The posterior distribution of last iteration (N=1000 particles) and the mean value is shown.

The separation between west African and MKK population occurred around 36 kyrs to 40 kyrs ago, also with a great amount of gene flow until present, with migration rate on magnitude of 10^−3^. The separation between CEU and GIH occurred around 36 kyrs to 38 kyrs ago, with ongoing migration on the magnitude of 10^−3^ until present. Comparisons with statistically-phased data suggest that the impact of phasing error on our PSMC-ABC method is less dramatic than for MSMC analysis, however when using haplotypes phased by SHAPEIT, a larger proportion of the genome coalesced ~50,000 years ago than when fosmid-phased haplotypes are used (Supplemental Figure 9, Supplemental Figure 10). This may result in larger amounts of inferred gene flow when using statistically phased data.

### Discussion

The utility of phase-resolved genome sequence data in the interpretation of variants impacting gene expression, transcription factor binding, human disease, and genome assembly has motivated the development of multiple approaches for determining phase. Here, we focus on samples phased using fosmid-based dilution haplotyping, and analyze a diverse set of eight phase-resolved human genomes. As expected, we find that phase results improve with increasing number of sequenced clones. We also demonstrate that statistical phasing performs well using existing reference panels, particularly when the panel captures population variation form the studied individuals. Nonetheless, the resulting phase-errors are sufficient to impact inference of population history using the MSMC model. We find that the statistically phased haplotypes show a more recent inferred population split time, perhaps due to phasing bias that make haplotypes appear more similar than they truly are. This effect is particularly noticeable for comparisons involving more deeply diverged population samples that are not well-phased using existing reference panels.

Existing PSMC and MSMC approaches represent important methodological advances and have had a clear impact on the inference of population history using individual genome sequences. However, these approaches provide only a qualitative sense of population separation history. Here, we describe the fitting of a standard Isolation with Migration model to cross-population TMRCA distributions inferred from PSMC. This allows the acquisition of parameter estimates under standard models widely utilized in population genetic inference. However, as expected, multiple combination of split time and migration rate are sometimes indistinguishable, highlighting the difficulty of inferring split times with the presence of migration (PRITCHARD 2011). This is partly due to the limitations of discretizing time and the poor resolution for recent history when given two haplotypes. Additionally, we find very high levels of migration for recent population splits (MKK and CEU, GIH and CEU, YRI and MKK), values which might be overestimated because of the high uncertainty for estimates of recent population history.

The split times inferred using our ABC method are generally concordant with the time when relative cross-coalescence rate dropped to 50% as inferred using MSMC, however our method provides a narrower range while quantifying the level of subsequent migration (Table 3). Utilizing this approach with fully phased haplotypes from nine populations, we provide additional estimates of key population separation in human population history. Overall, our estimates are broadly consistent with other contemporary methods (Supplemental Table 6) and our estimates are consistent with the timing of the most recent common ancestor of African and non-African mitochondrial DNA, around 78,300 years ago and the timing of the mitochondrial MRCA for all modern humans at 157,000 years ago (FU *et al.* 2013).

**Table 3.** **Posterior estimates of split time and migration rate using IM model**. We report the mean, median and 95% credible intervals for the posterior distribution. Migration rate are in log10 scale. We set migration continuing to the present for recent separations.

|  | Split Time (in kyrs) |  |  |  | MigrationRate (in log<sub>10</sub> scale) |  |  |  | Migration End (in kyrs) |  |  |  |
| --- | --- | --- | --- | --- | --- | --- | --- | --- | --- | --- | --- | --- |
|  | Mean | Median | 95% lower HPD | 95% higher HPD | Mean | Median | 95% lower HPD | 95% higher HPD | Mean | Median | 95% lower HPD | 95% higher HPD |
| YRI-CEU | 73.3 | 72.5 | 70.2 | 81.6 | -3.71 | -3.70 | -4.12 | -3.41 | 27.2 | 30.1 | 6.2 | 38.7 |
| MKK-CEU | 53.9 | 53.9 | 52.9 | 55.1 | -2.34 | -2.22 | -2.66 | -2.04 | - | - | - | - |
| GIH-CEU | 37.2 | 37.2 | 36.2 | 38.2 | -2.88 | -2.87 | -3.17 | -2.60 | - | - | - | - |
| San-CEU | 129.5 | 128.8 | 121.3 | 140.9 | -3.95 | -3.96 | -4.07 | -3.83 | 37.2 | 37.1 | 33.4 | 41.5 |
| Mbuti-CEU | 117.6 | 116.9 | 103.1 | 139.1 | -3.73 | -3.73 | -3.82 | -3.63 | 34.6 | 34.4 | 30.6 | 39.1 |
| YRI-MKK | 38.2 | 38.1 | 36.2 | 40.6 | -2.15 | -2.16 | -2.30 | -2.00 | - | - | - | - |

Similar to previous results (SCHIFFELS and DURBIN 2014), the separation history between CEU and MKK populations was different from that observed between CEU and LWK (Luhya, another east African population). Two pulses of admixture have been estimated in the ancestors of the MKK, occurring 8 and 88 generations ago ((PAGANI *et al.* 2012; PICKRELL *et al.* 2014)). Since the impact of long segments of shared ancestry due to recent admixture is unclear, we masked out regions of recent European ancestry in our MKK sample using RFMix (MAPLES *et al.* 2013) (Supplemental Figure 11) and found that the MSMC curves are not altered when recent segments of European ancestry are masked (Supplemental Figure 12). Although such ancestral masking becomes increasingly imperfect for older admixture events, this suggests that long segments of shared ancestry due to recent admixture do not explain the latter divergence of Massai population compared to other African populations and supports a more complex ancient history for the Massai.

When constructing global haplotypes for individuals without trio phasing data available, we applied Prism to statistically link blocks together. Prism was designed to link much shorter phased segments into longer blocks. When applied to our phased haplotype blocks, we found that around 40% of blocks were assigned incorrectly, resulting in switch errors every 2 Mbp. However, we found very similar MSMC curve using global haplotypes constructed by Prism with those constructed with trio phasing data (Supplemental Figure 13), indicating long switch errors have little effect on such inference. This is reassuring since we are using Prism to construct global haplotypes for four individuals; but, the inferred split times involving the San and Mbuti populations are still likely underestimated.

Our results indicate that the separation of the studied human populations was a gradual event, with substantial genetic exchange continuing after an initial split, a finding consistent with hypotheses of long-standing ancient population structure in Africa (reviewed in (HARDING and MCVEAN 2004; HENN *et al.* 2012)). We provide a comparison of PSMC and MSMC based methods with other contemporary methods on inferring population separation history and our results emphasize the importance of accurately phased haplotypes on MSMC analyses, especially for more ancient splits.

## Acknowledgments

We thank Jacob Kitzman, Peedikayil Thomas, Jeffrey W. Innis, and the University of Michigan DNA Sequencing Core Facility for guidance on fosmid pool construction and sequencing.

## Web resources

The reconstructed haplotypes are available for download from DataDyrad under accessions XXX.

---

**SFigure 1.**
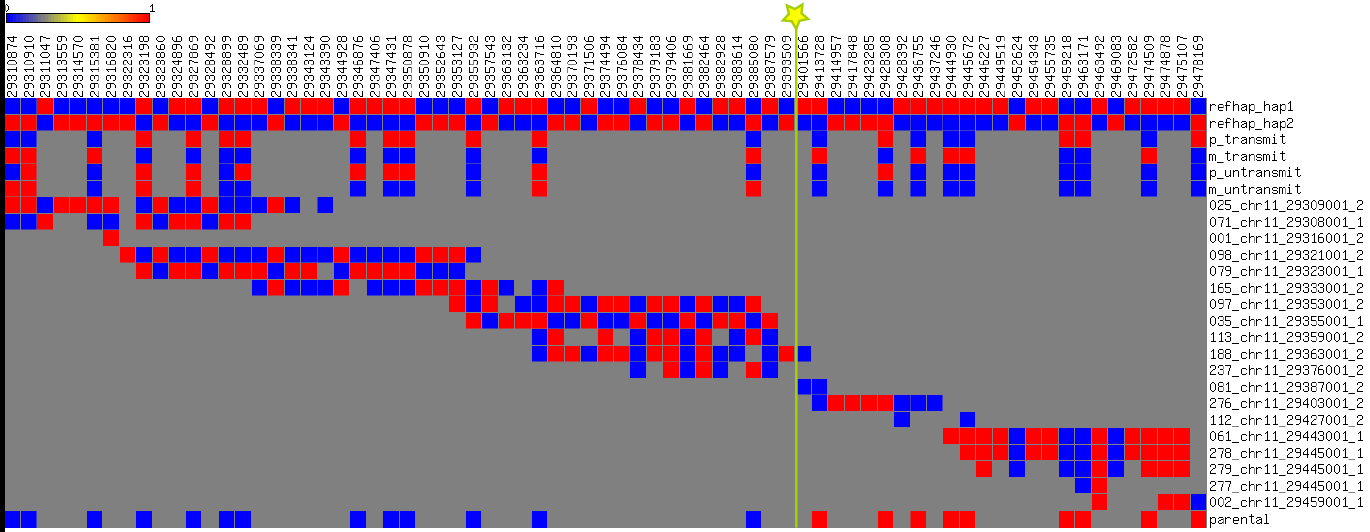
**Illustration of ReFHap‘s phasing result and a switch error**. Each column corresponds to a SNP position, with blue indicating the reference allele and red the alternative. The first two rows are the haplotype prediction by ReFHap, followed by four rows showing HapMap phase based on trio transmission. This is followed by 12 rows depicting clone genotypes. The last row indicates the parental allele assigned for RefHap haplotype based on HapMap phasing. In the last row, blue indicates paternal allele and red indicates maternal allele. The line with a star shows where the switch error occurred.

**SFigure 2.**
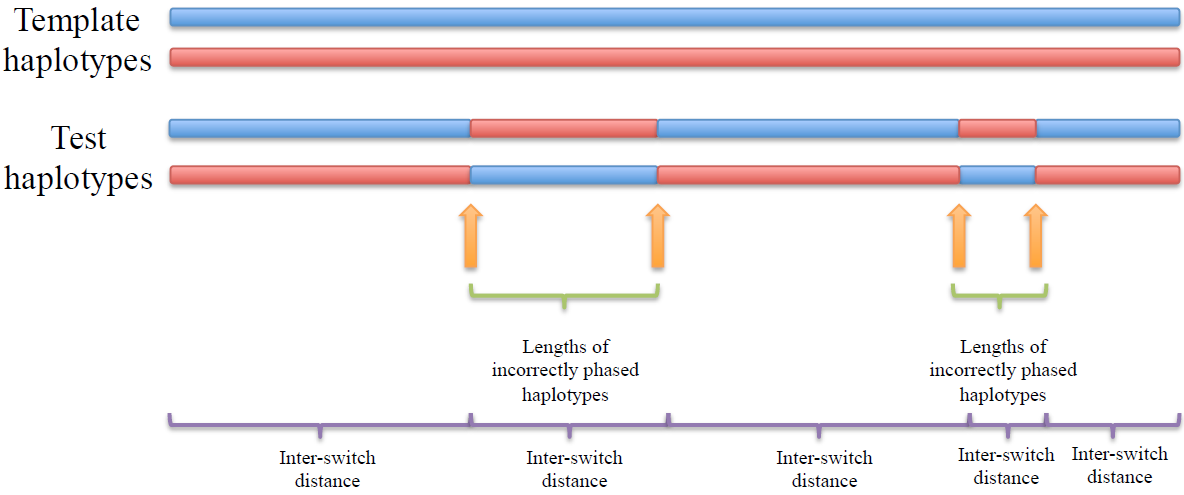
**Illustration of the metrics used to quantify phasing errors**. We illustrate switch error (green bracket), inter-switch distance (purple bracket) and length of incorrectly phased haplotypes(green bracket) when comparing test haplotypes with template haplotypes.

**SFigure 3.**
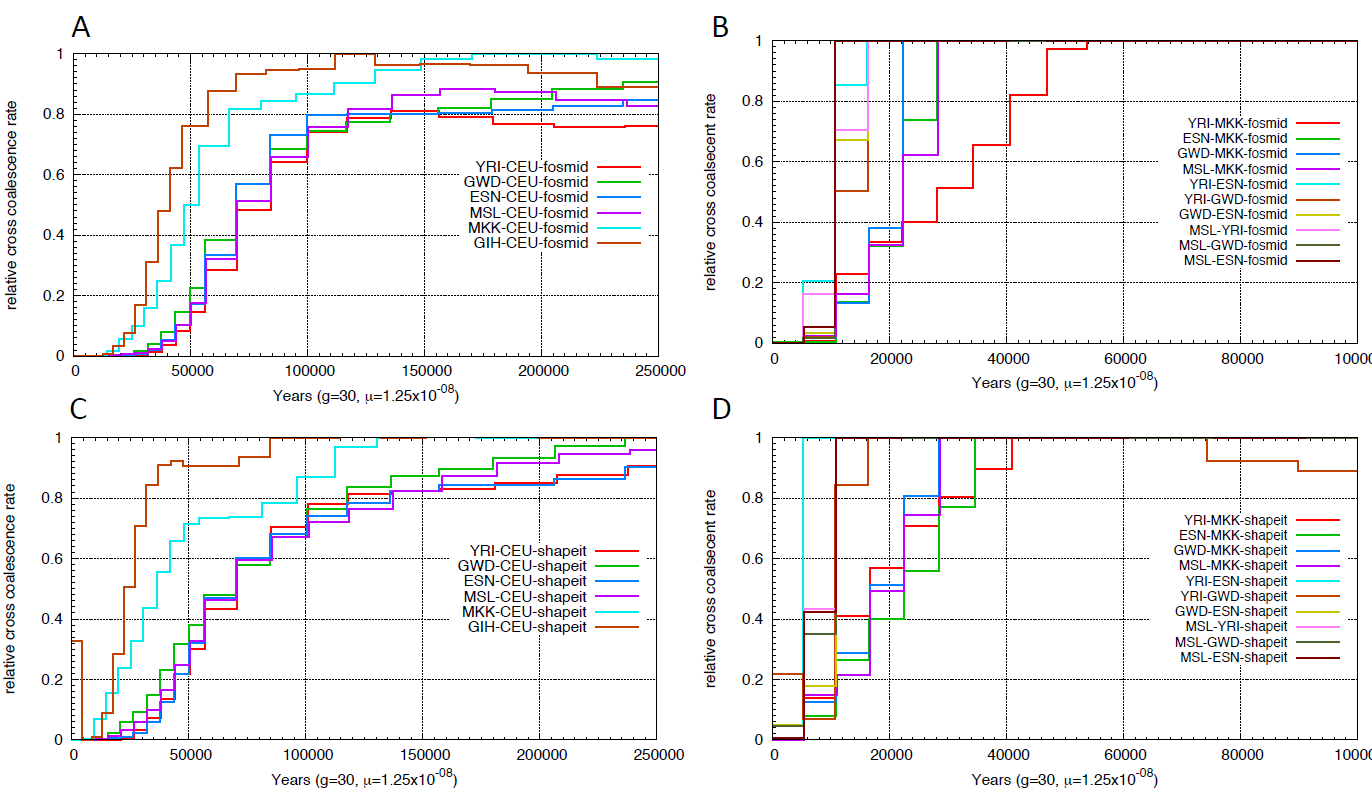
**Relative cross coalescence rate inferred using MSMC**. We applied msmc on four haplotypes, two from each population. We compared the relative cross coalescence curve using physically phased haplotypes (A,B,E,F) vs haplotypes phased using SHAPEIT with 1000 Genomes Phase1 panel (C,D,G,H).

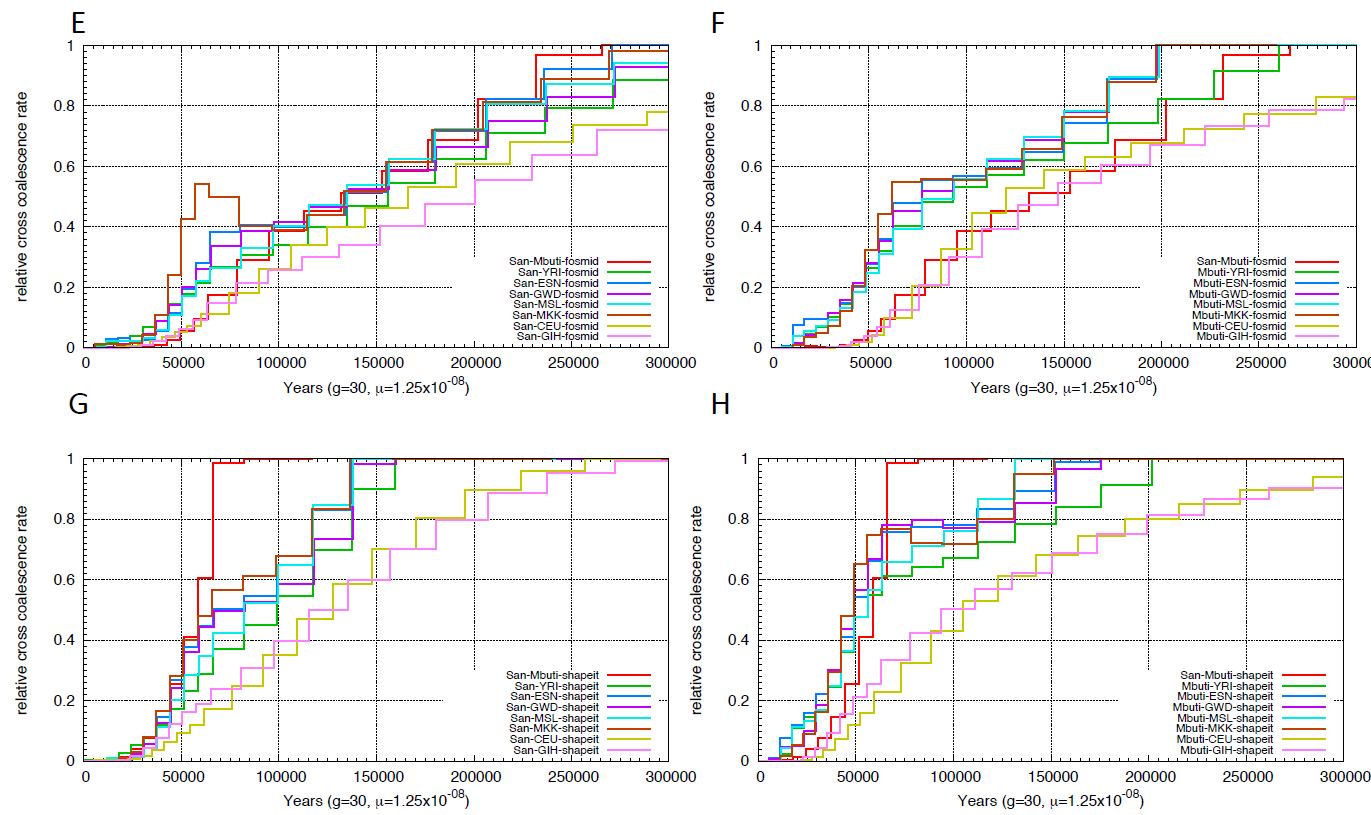

**SFigure 4.**
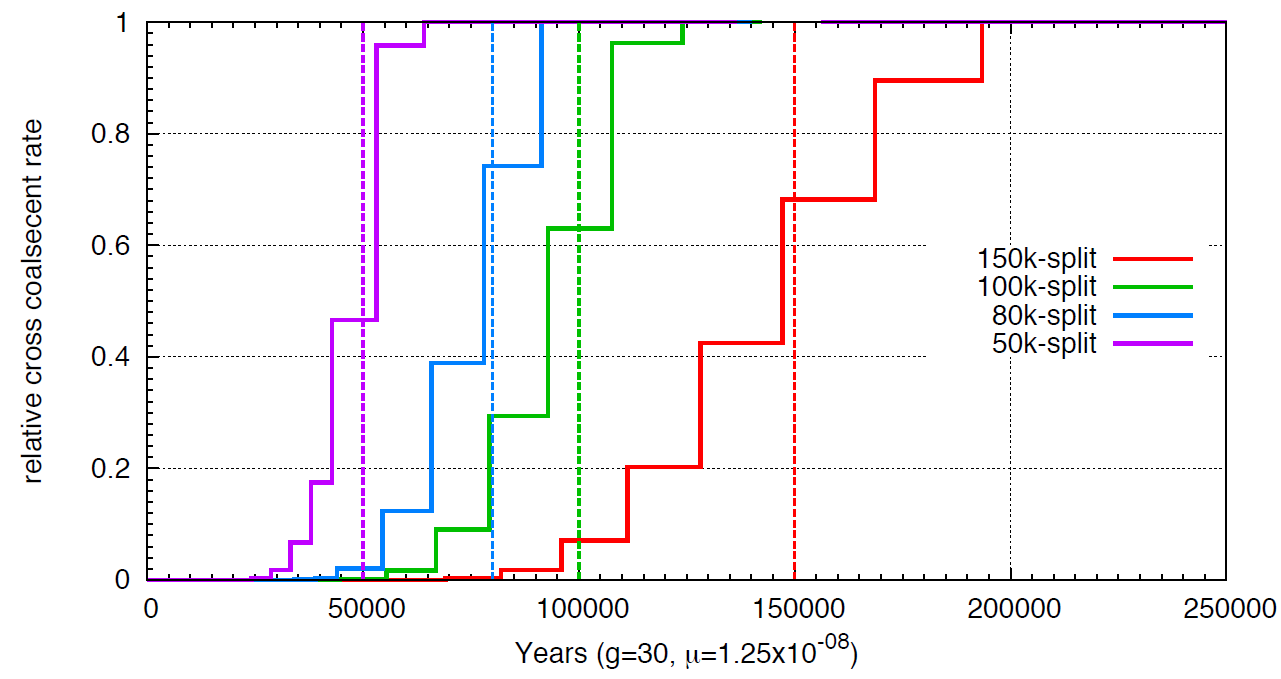
**Relative cross coalescence rate inferred using MSMC on simulated data**. We performed simulation using MaCS (100 30M sequences for each individual) with a clean population split at 50 kyrs, 80 kyrs, 100 kyrs and 150 kyrs ago. We applied msmc on simulated sequences and plotted the relative cross coalescence curve.

**SFigure 5.**
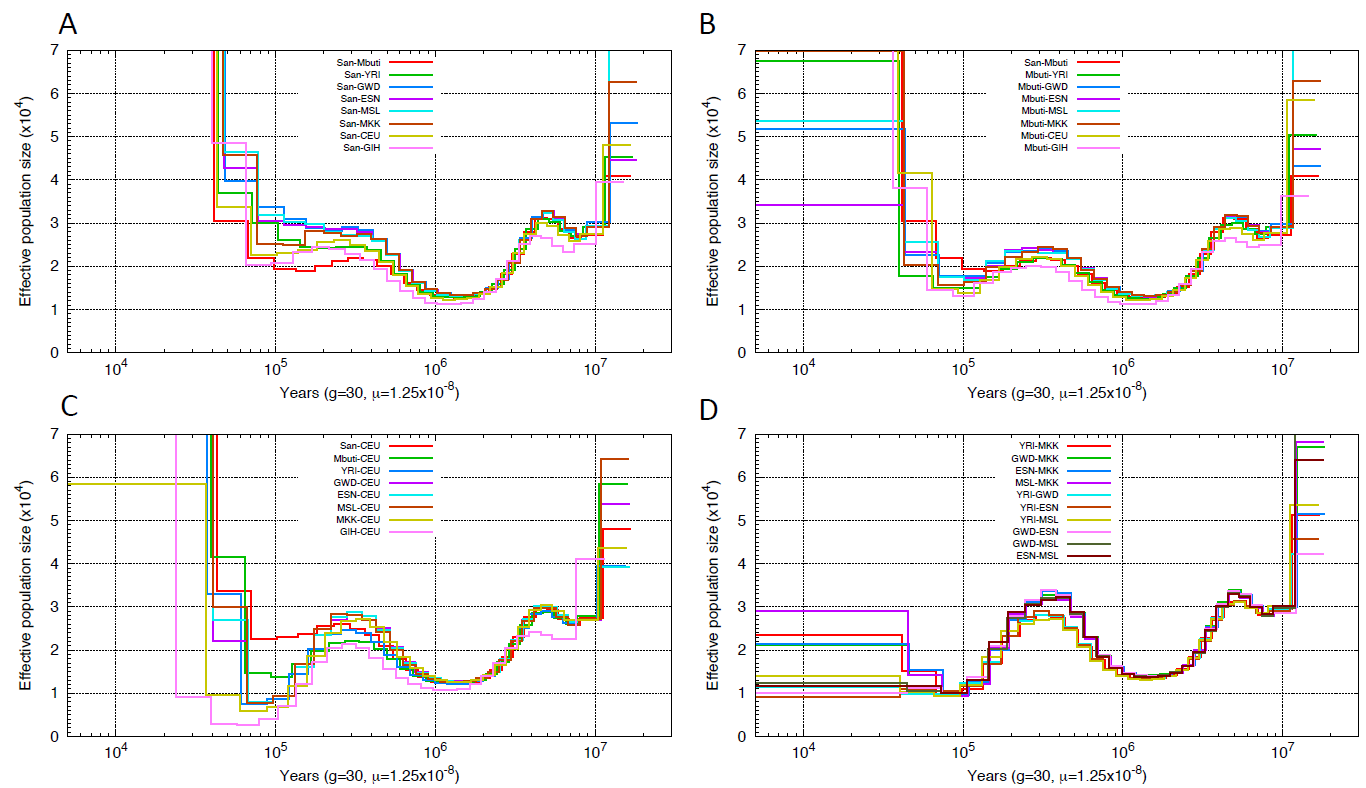
**PSMC on pseudo-diploid genomes**. Population sizes inferred from combined autosomes, one haplotype from each population are shown. Sizes are the average from 4 haplotype configuration, namely hap1-hap1,hap1-hap2,hap2-hap1,hap2-hap2. Haplotypes are constructed using fosmid pool approach.

**SFigure 6.**
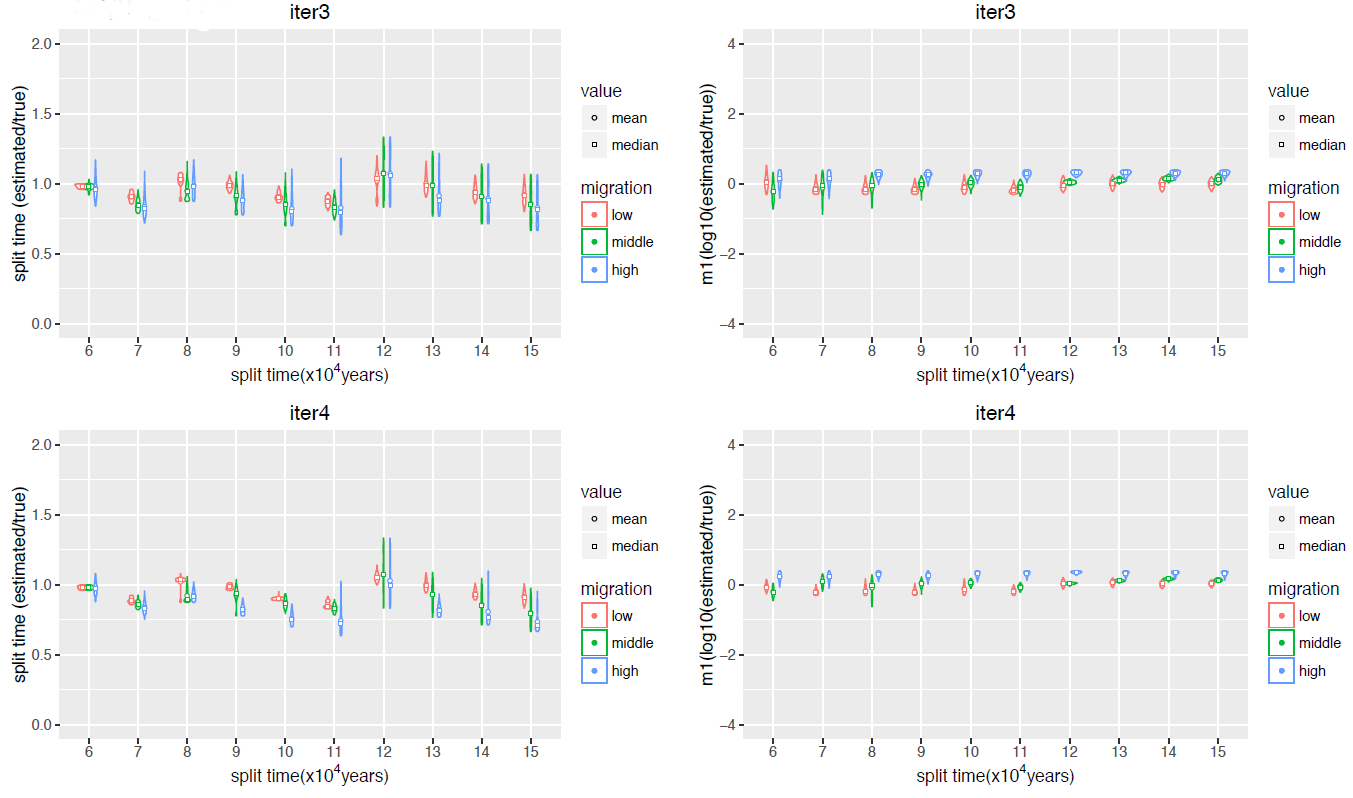
**Simulation results on inferring split time and migration using the combined approach of PSMC and ABC**. We tested our approach using simulated data of two individuals representing African and European population and having a split time from 60 kyrs to 150 kyrs ago, with subsequent migration until 30 kyrs ago. We tested three level of migration rate, 2×10^−5^ (low), 10×10^−5^ (middle), 40×10^−5^(high) and plotted the posterior distribution, mean and median value of split time (estimated/true) and migration rate (log10(estimated/true)) of iteration3 and iteration4 of our ABC approach.

**SFigure 7.**
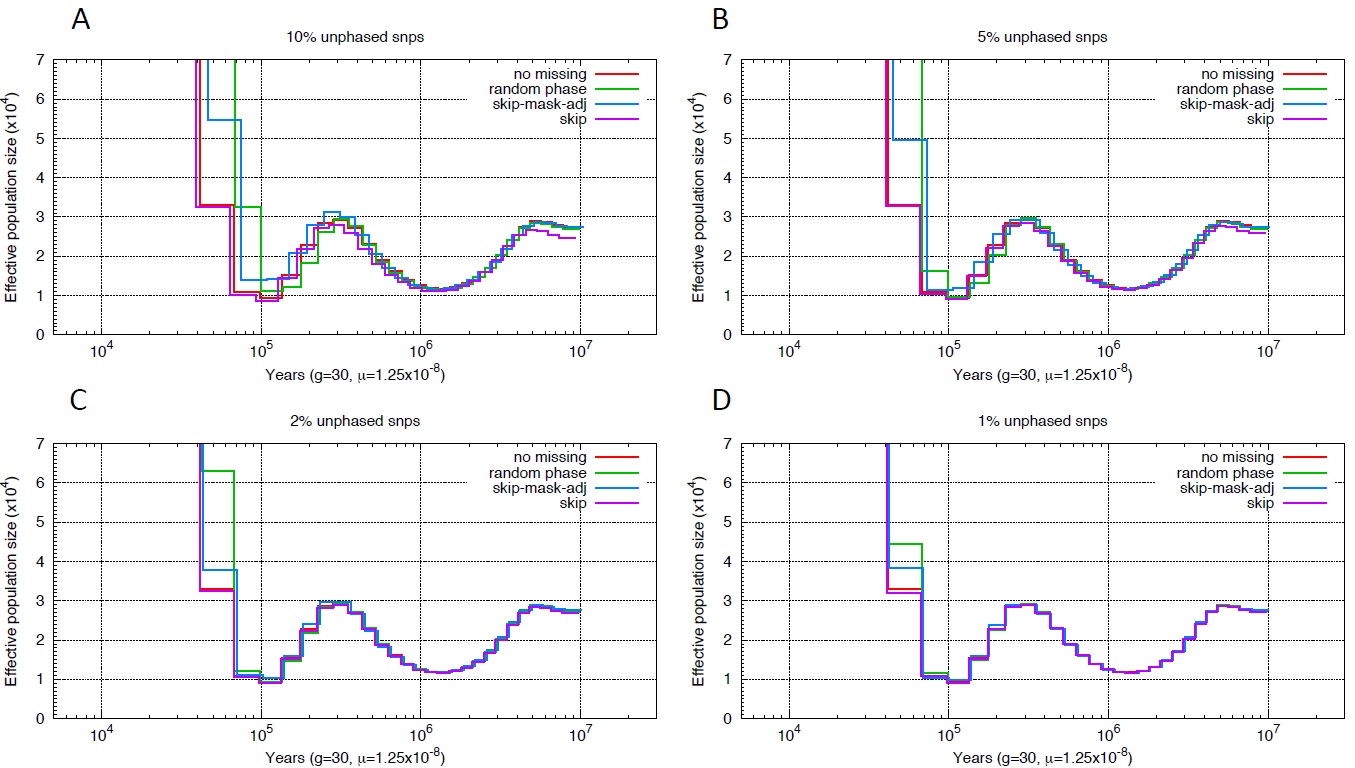
**Simulation results on different approaches to deal with unphased SNPs**. We simulated sequences with different levels of unphased snps (1%, 2%, 5%, 10%) and evaluated three different methods to deal with unphased snps, 1) randomly assigning the phase (green lines), 2) marking unphased snps as missing data and removing all blocks of homozygous calls that ended in an unphased heterozygous site (blue) and 3) merely marking unphased snps as missing data (purple).

**SFigure 8.**
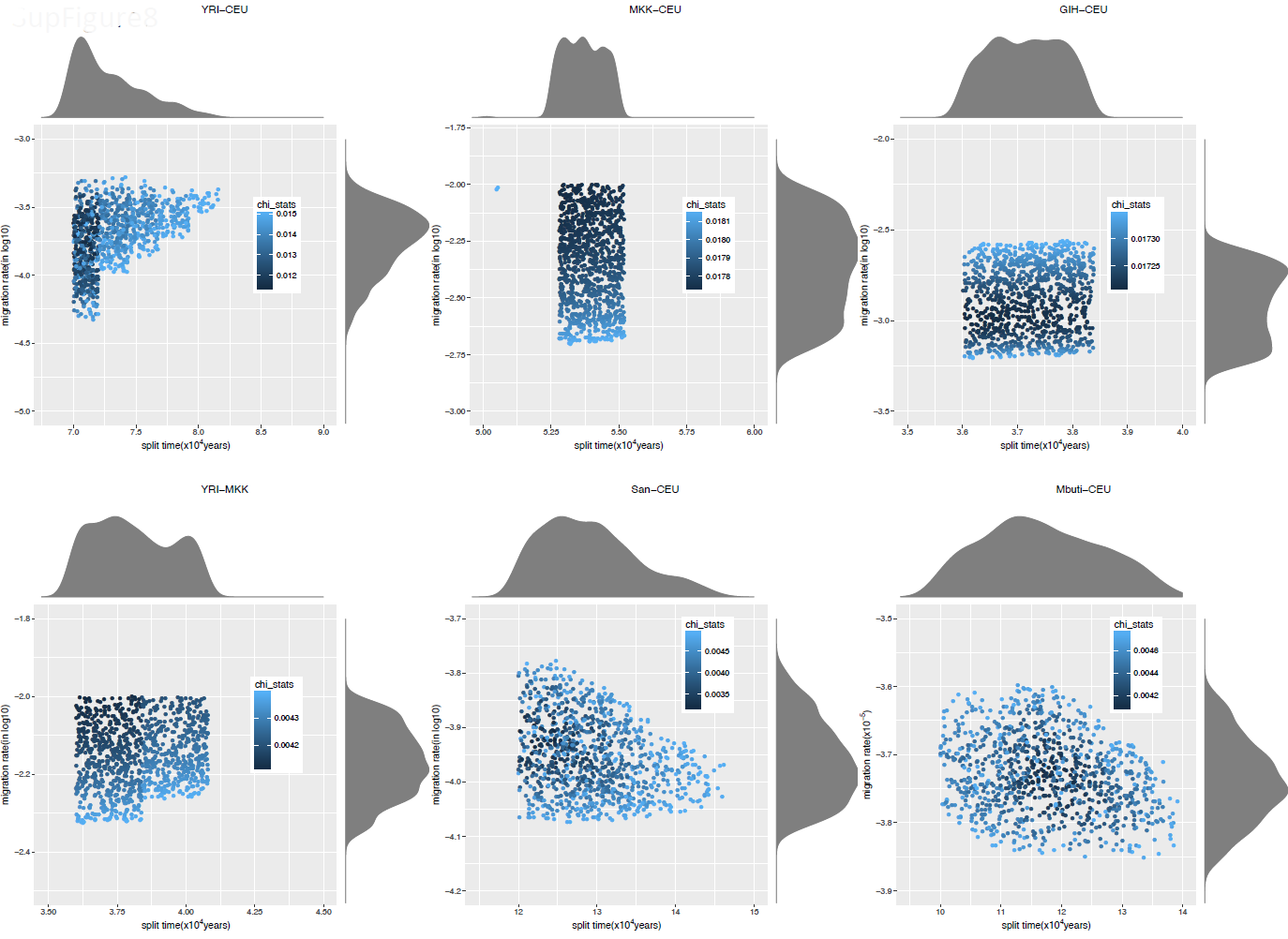
**Posterior distribution of split time and migration rate inferred using ABC**. We applied ABC-SMC to infer split time and migration rate based on the inferred TMRCA distribution obtained from PSMC. For each pair of populations, we plotted the posterior distribution of split time and migration rate. The color represents chi square distance between the TMRCA distribution from the observed data and the model.

**SFigure 9.**
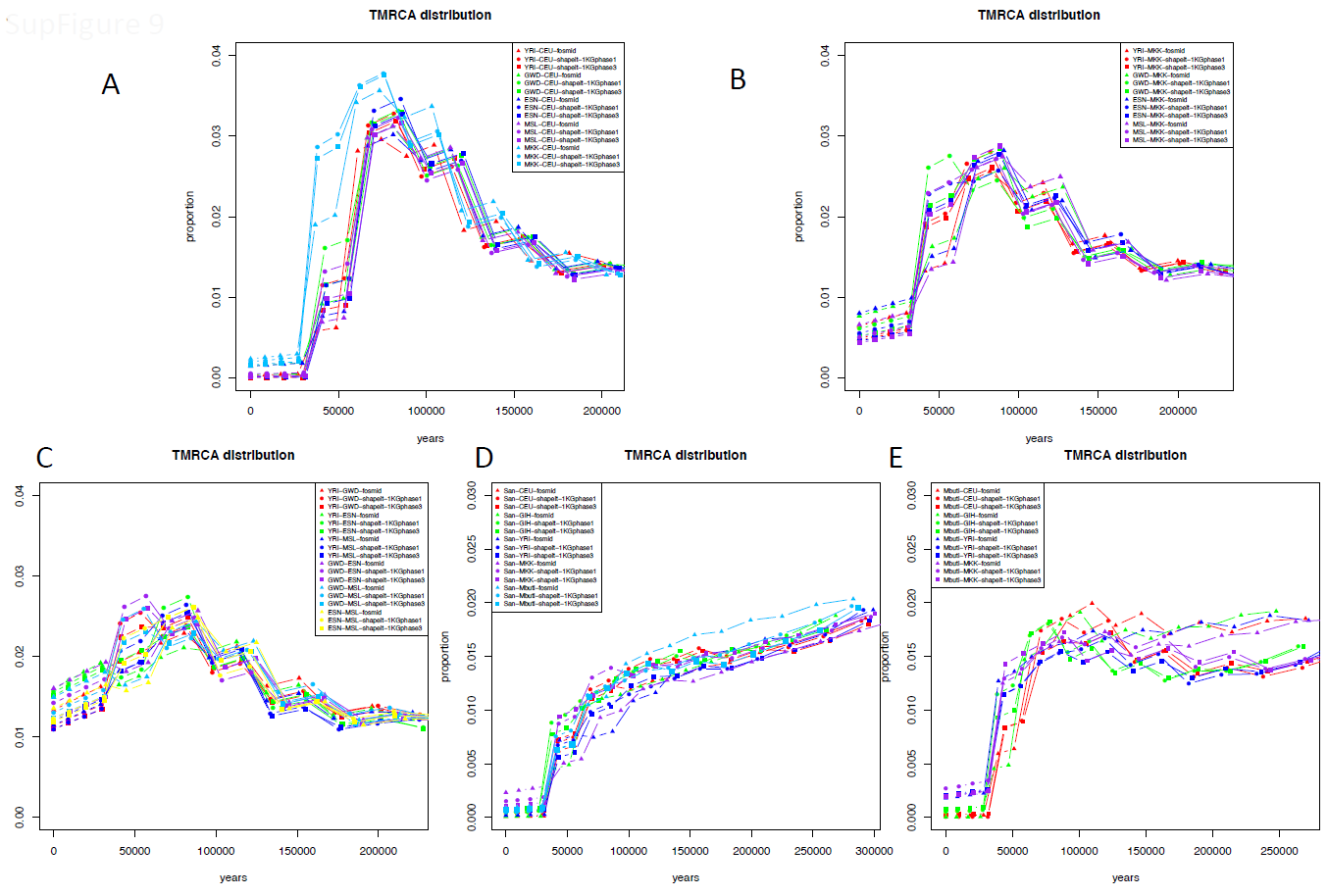
**TMRCA distribution inferred using PSMC**. The figure shows the left tail of TMRCA distribution inferred using PSMC on pseudo-diploid individuals for comparisons involving CEU (A), MKK (B), GWD (C), San (D), and Mbuti (E). Each plot shows the TMRCA distribution inferred using haplotypes phased using fosmid data (triangle) and phased using SHAPEIT with 1000 Genomes Phase1 (circle) and Phase3 (square) reference panels.

**SFigure 10.**
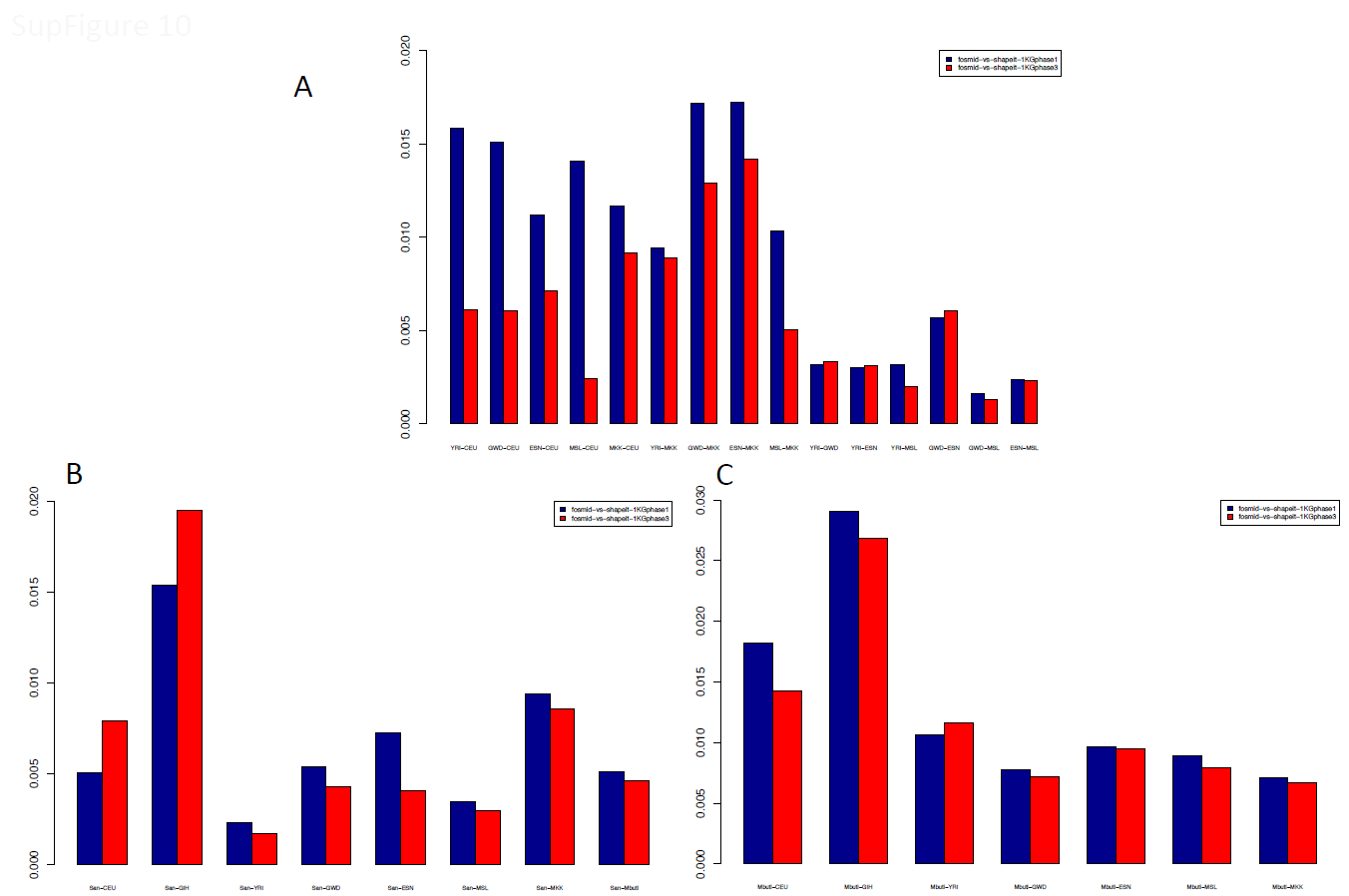
**Chi-square distance between TMRCA distributions using different haplotypes**. We plotted the chi-square distance between TMRCA distributions obtained using different haplotypes phased using fosmid data and phased using SHAPEIT with 1000 Genomes Phase1 (blue) and Phase3 (red) reference panels.

**SFigure 11.**
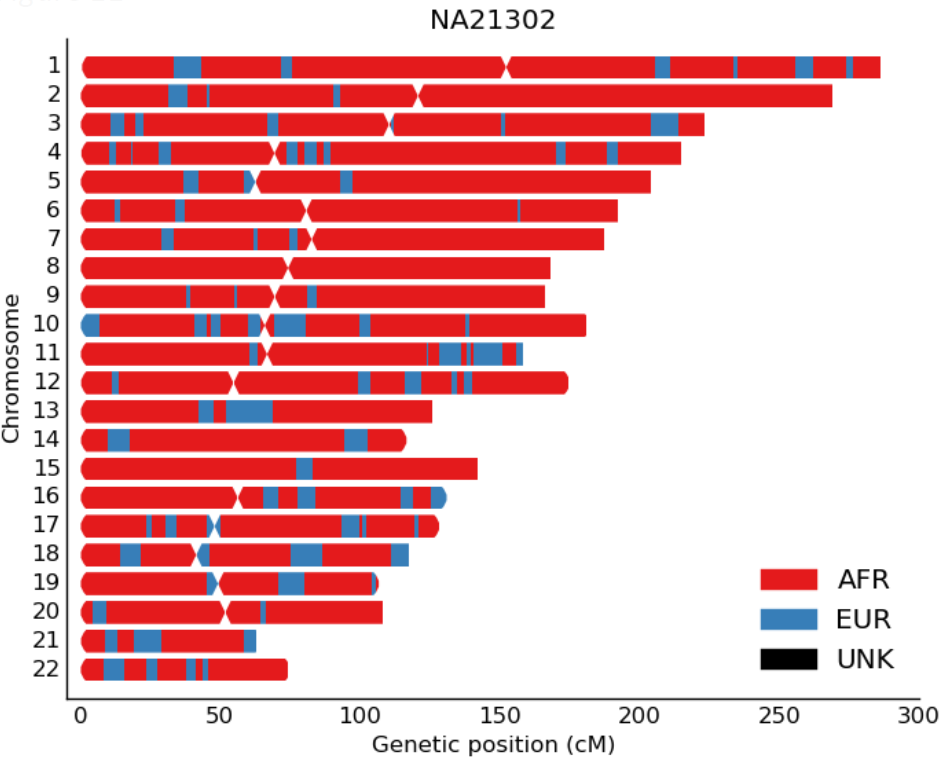
**Recent European ancestry inferred by RFMix in Massai individual NA21302**. The genomic locations of European ancestry (colored blue) in Massai individual NA21302 are shown

**SFigure 12.**
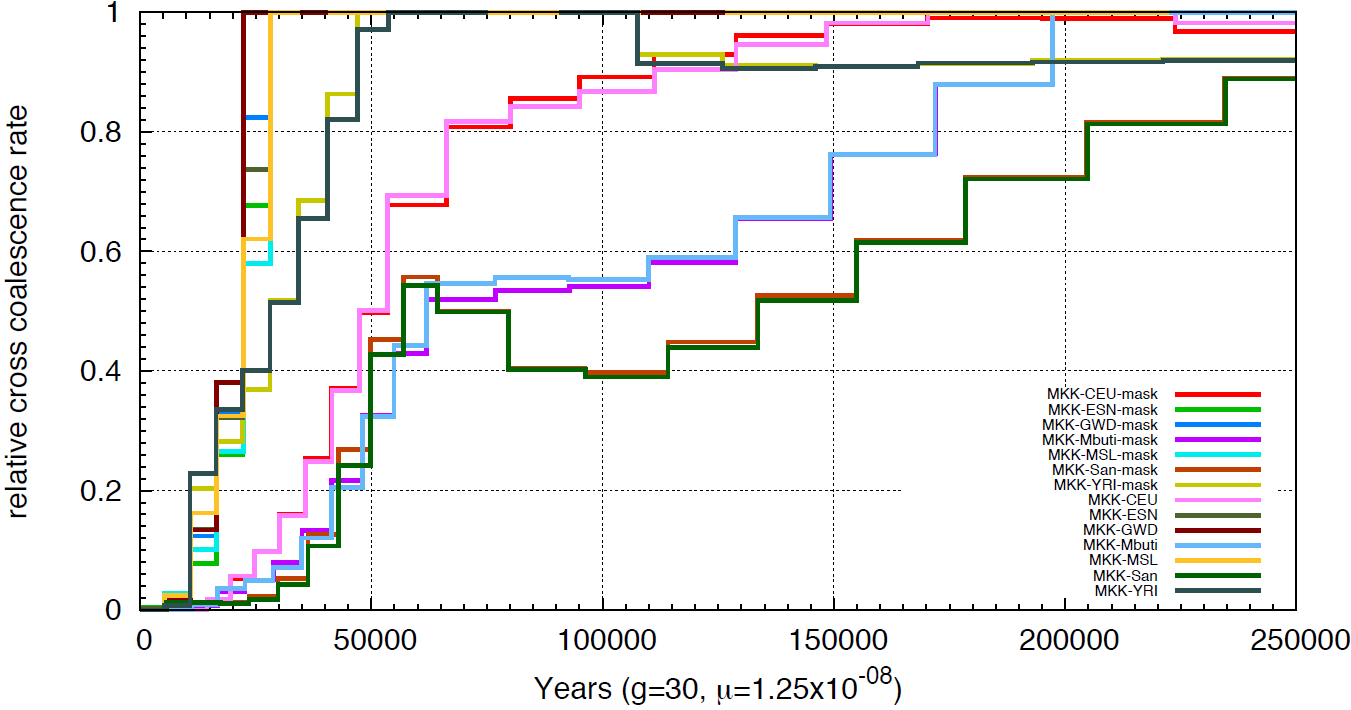
**Relative cross coalescence rate inferred using MSMC with and without masking out European ancestry from Massai individual**. We applied MSMC on MKK and every other population with and without masking European ancestry from the Massai individual.

**SFigure 13.**
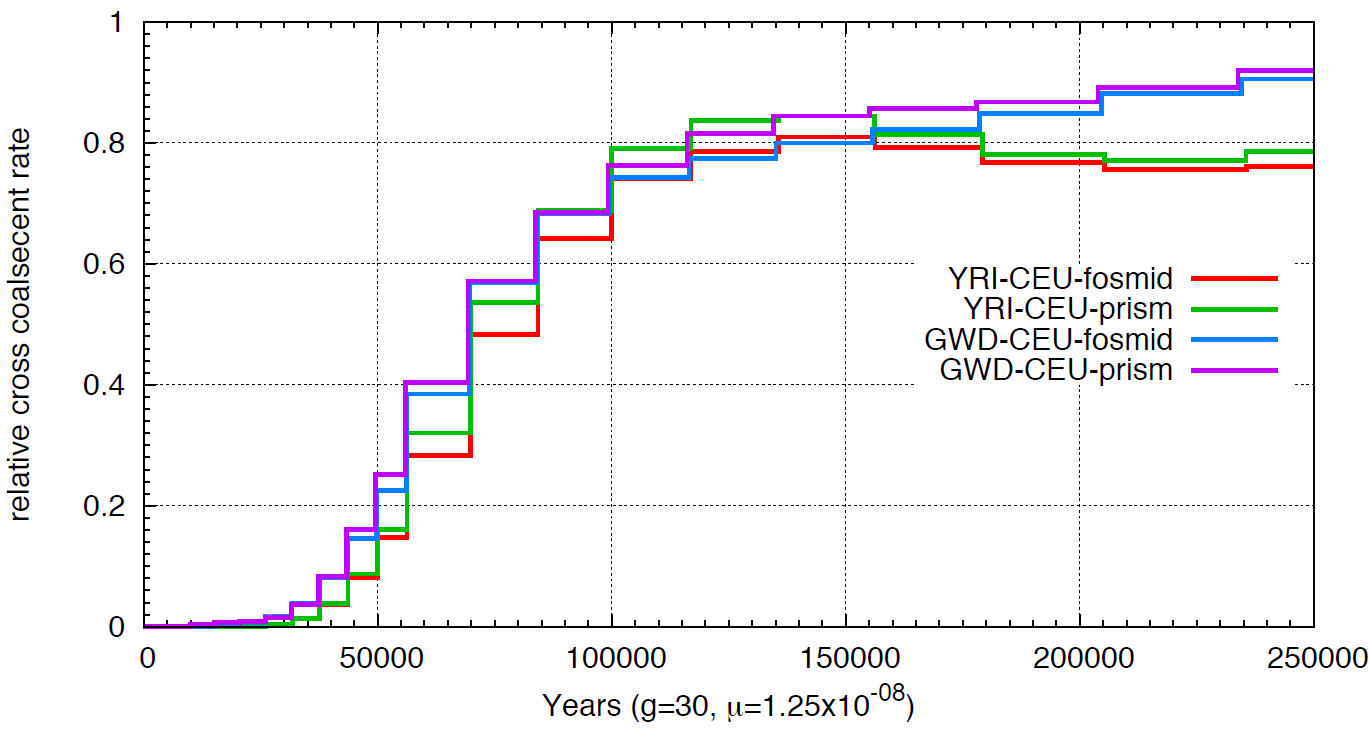
**Comparison of relative cross coalescence rate inferred using MSMC on fosmid haplotypes constructed by trio phasing data or by Prism**. We compared the relative cross coalescence curve for YRI-CEU and GWD-CEU, where the global haplotypes
1.for YRI and GWD are either constructed by trio phasing data or by Prism.

**STable 1.**
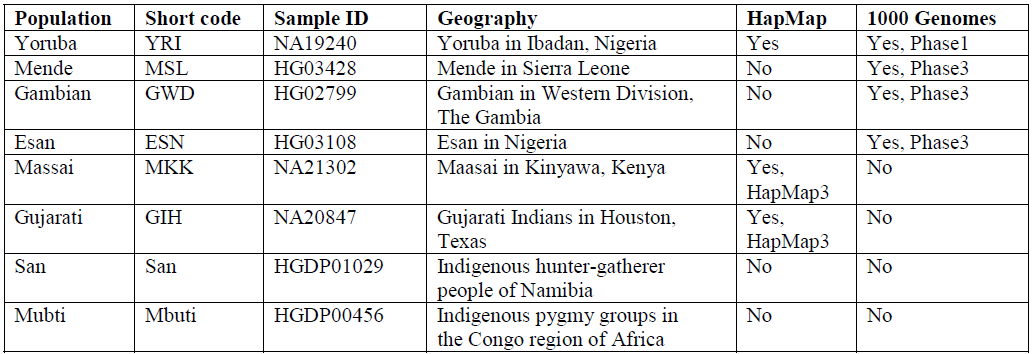
Summary of population geographic information and presence in HapMap or 1000 Genomes Project.

**STable 2.**
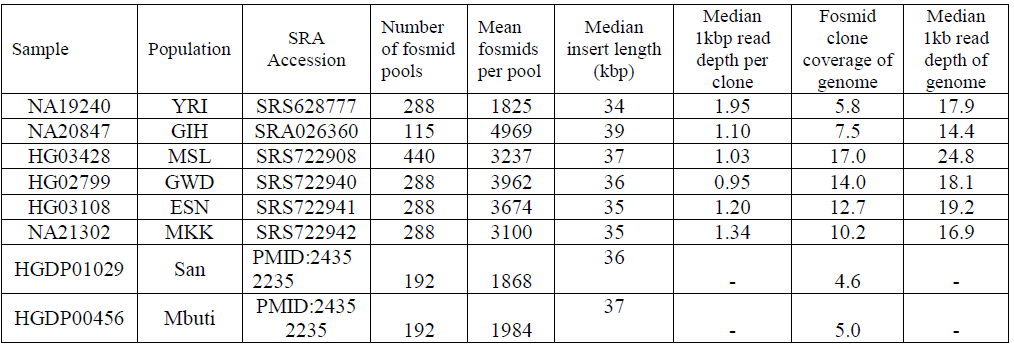
Summary of clone statistics of fosmid pool sequencing.

**STable 3.**
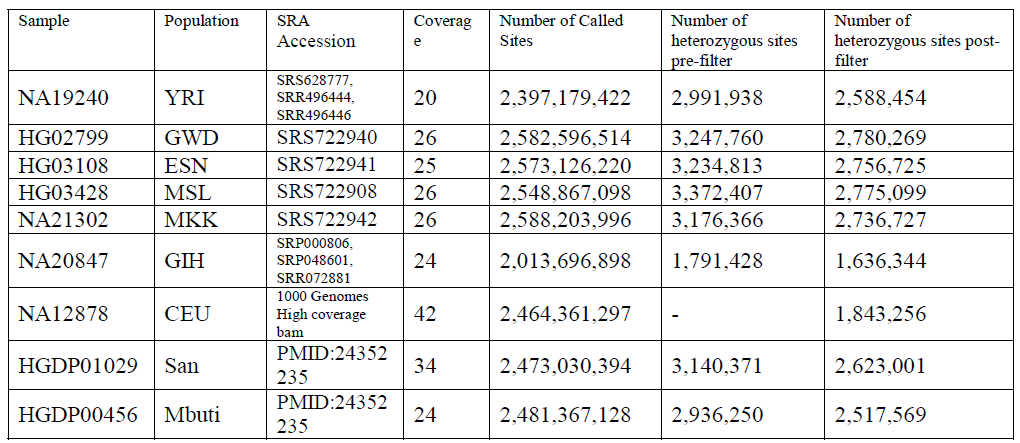
Summary of variant calling for whole genome sequencing.

**STable 4.**
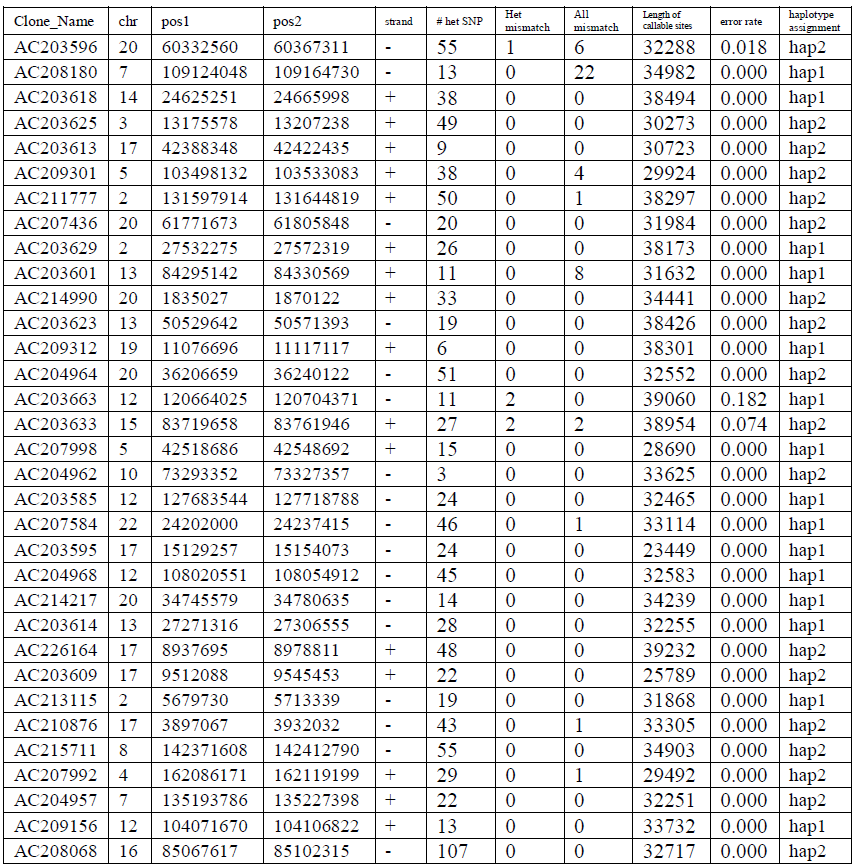
**Comparison of fosmid-resolved haplotype for NA19240 with Sanger sequenced fosmid clones**. Comparison of our haplotypes with the sequence of 33 fosmid clones from the same individual that were previously sequenced using standard capillary sequencing (hap1 refers to paternal allele, hap2 refers to maternal allele). A total of 5 out of 1013 heterozygous SNP error occurred when assigning 33 clones into haplotype, an error rate of 0.5%. substitution errors out of 1,102,213 bp total sequences yielded a sequence error rate of 0.005%.

**STable 5.**
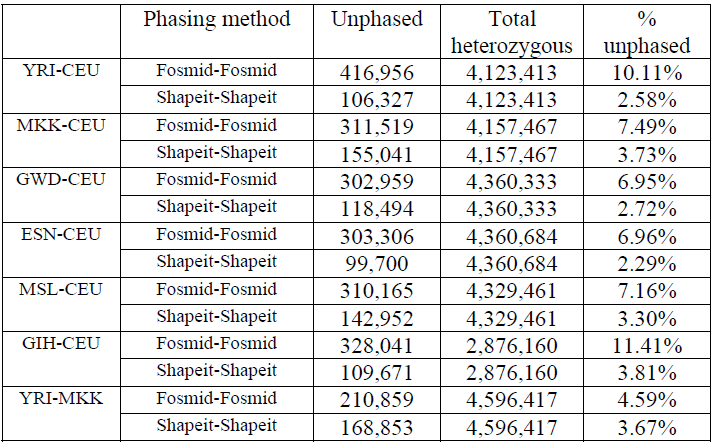
**Percentage of unphased SNPs**. We summarized the proportion of unphased SNPs for each population combination. For ‘Shapeit’ we refer to applying SHAPEIT with 1000 Genomes Phase I reference panel.

**STable 6.**
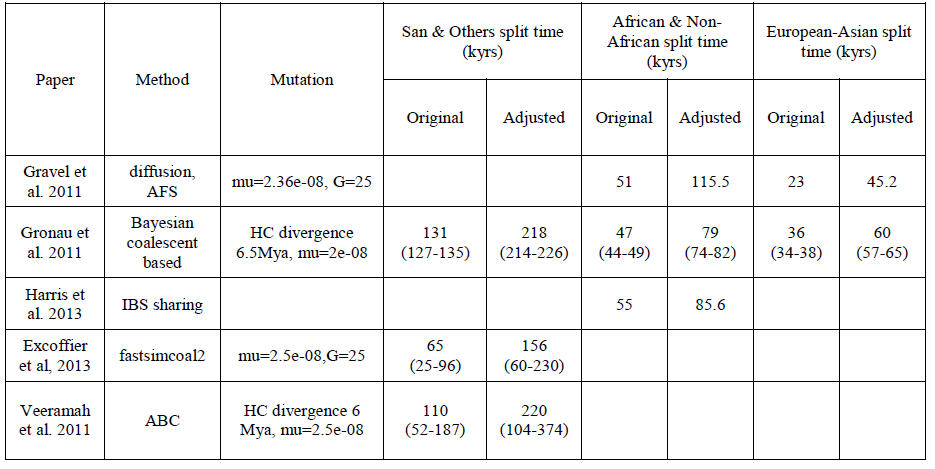
**Split time estimation from previous studies**. Reported estimates are adjusted by using the same mutation rate 1.25*10^−8^ bp/generation and generation time 30 years.

